# Point-of-Use Detection of Environmental Fluoride via a Cell-Free Riboswitch-Based Biosensor

**DOI:** 10.1101/712844

**Authors:** Walter Thavarajah, Adam D. Silverman, Matthew S. Verosloff, Nancy Kelley-Loughnane, Michael C. Jewett, Julius B. Lucks

**Affiliations:** Department of Chemical and Biological Engineering, Northwestern University, 2145 Sheridan Rd, Evanston, IL, 60208, USA; Interdisciplinary Biological Sciences Graduate Program, Northwestern University, 2204 Tech Drive, Evanston, IL, 60208, USA; Center for Synthetic Biology, Northwestern University, 2145 Sheridan Rd, Evanston, IL, 60208, USA; Center for Water Research, Northwestern University, 2145 Sheridan Rd, Evanston, IL, 60208, USA; Materials and Manufacturing Directorate, Air Force Research Laboratory, Wright-Patterson Air Force Base, Ohio 45433, United States

## Abstract

Advances in biosensor engineering have enabled the design of programmable molecular systems to detect a range of pathogens, nucleic acids, and chemicals. Here, we engineer and field-test a biosensor for fluoride, a major groundwater contaminant of global concern. The sensor consists of a cell-free system containing a DNA template that encodes a fluoride-responsive riboswitch regulating genes that produce a fluorescent or colorimetric output. Individual reactions can be lyophilized for long-term storage and detect fluoride at levels above 2 parts per million, the EPA’s most stringent regulatory standard, in both laboratory and field conditions. Through onsite detection of fluoride in a real-world water source, this work provides a critical proof-of-principle for the future engineering of riboswitches and other biosensors to address challenges for global health and the environment.

## Introduction

Safe drinking water availability is an important contributor to public welfare (*1*). However, safe water sources are not available to a large portion of the globe, with an estimated 3 billion people using water from either an unsafe source or a source with significant sanitary risks (*2*). One particularly dangerous contaminant is fluoride, which leaches into groundwater from natural sources. Long-term exposure to fluoride concentrations above 2 parts per million (ppm) can cause dental and skeletal fluorosis, heavily burdening communities in resource-limited settings (*3*). Though large-scale remediation strategies are available, they are resource-intensive and difficult to deploy (*3, 4*). This problem is compounded by the reliance of gold-standard sensing methods on expensive analytical equipment, making detection difficult in areas with the greatest need (*4*). To facilitate targeted remediation and empower affected individuals, there is a pressing need for a more practical, rapid, and field-deployable solution to monitor the presence of fluoride in water.

Because they can be freeze-dried and rehydrated on-demand, cell-free systems have recently become a promising platform for field-deployable molecular diagnostics (*5, 6*). These systems typically consist of cellular gene expression machinery along with the required buffers, energy sources and co-factors necessary to support gene expression from added DNA templates (*7*). Unlike whole cells, cell-free platforms offer an open, easily tunable reaction environment, expediting the design process for genetically encoded programs (*7*). Furthermore, they circumvent the analyte toxicity, host mutation, and biocontainment concerns limiting cellular sensors (*8*).

We sought to leverage the advantages of cell-free biosensing platforms to create a new approach for monitoring for the presence of fluoride in water using a fluoride-responsive riboswitch that regulates the expression of the crcB fluoride efflux pump in *Bacillus cereus* (*7, 9, 10*). By configuring the *B. cereus* crcB fluoride riboswitch to control the transcription of downstream reporter genes (*11*), we show that a cell-free gene expression system can activate both protein and RNA reporter expression in the presence of fluoride. With an enzymatic colorimetric reporter, we demonstrate detection of fluoride concentrations at the Environmental Protection Agency (EPA) Secondary Maximum Contaminant Level of 2 ppm (*12*). We also demonstrate that our fluoride biosensor can be lyophilized for long-term storage and distribution, allowing us to detect fluoride in unprocessed groundwater obtained and tested onsite in Costa Rica. This work exemplifies the potential of riboswitches as practical biosensing tools and lays the foundation for utilizing cell-free biosensing systems in rapid and field-deployable water quality diagnostics to address pressing challenges in global health.

## Results

### Fluoride riboswitch control of reporter expression in cell-free reactions

Our point-of-use diagnostic consists of a cell-free system containing a fluoride biosensor DNA template that can be freeze-dried and stored. Rehydration activates the biosensor, which encodes the fluoride riboswitch and a reporter gene that produces a detectable output if fluoride is present (**Figure 1A, B**). As a starting point, we sought to characterize the regulatory activity of the *B. cereus* crcB riboswitch in the cell-free reaction environment. Previous characterization of the riboswitch’s cotranscriptional folding mechanism (**Figure 1B**) confirmed that it functions with *E. coli* RNA polymerase (*11*), allowing us to use it in *E. coli* cell-free extract. We therefore constructed a reporter plasmid containing the riboswitch sequence followed by a strong ribosome binding site (RBS) and the coding sequence of the reporter protein superfolder green fluorescent protein (sfGFP), all placed downstream of a constitutive *E. coli* σ70 promoter (all plasmid details in **Supplemental Table 1**).

**Figure 1.**
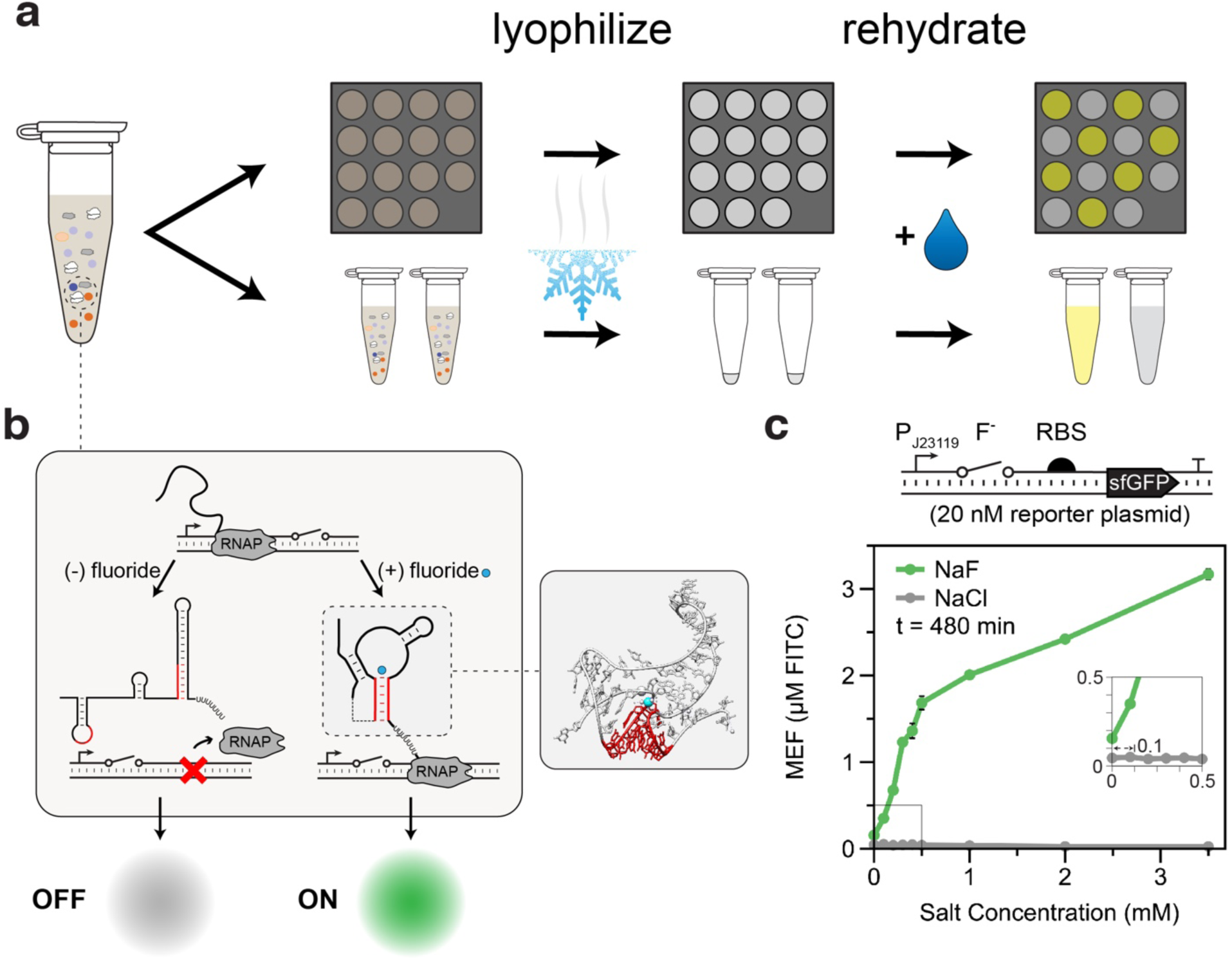
Cell-free fluoride biosensor engineering strategy. **(a)** Schematic for lyophilization of a cell-free reaction in tubes or on paper disks. Rehydration with a water sample allows the designed biosensing reaction to proceed to yield a detectable signal. **(b)** Schematic for fluoride riboswitch-mediated transcriptional regulation in cell-free extract. The riboswitch folds cotranscriptionally into one of two mutually exclusive states, depending on the presence of fluoride. In the absence of fluoride, the riboswitch folds into a terminating hairpin, precluding downstream gene expression. Fluoride binding stabilizes a pseudoknot structure (red paired region, inset from PDB:4ENC) that sequesters the terminator and enables the expression of downstream reporter genes. **(c)** Schematic of a cell free fluoride biosensor, consisting of a DNA template encoding the fluoride riboswitch controlling the expression of sfGFP. Eight-hour endpoint fluorescence measurements for reactions containing NaF (dark green) or NaCl (gray) are shown below. Error bars represent one standard deviation from three technical replicates.

After optimizing reaction conditions for riboswitch performance (**Supplemental Figure 1**), we determined the fluoride sensor’s dose-response to fluoride by titrating across a range of NaF concentrations. All tested conditions caused a measurable increase in expression over the OFF state, with induction seen at NaF concentrations as low as 0.1 mM (**Figure 1C**, green line and inset). This threshold is important, since 0.1 mM NaF is equivalent to the EPA’s 2 ppm secondary maximum contaminant level for fluoride in drinking water, its most stringent risk threshold (*12*). Importantly, the observed leak in the absence of NaF was also observed to be near zero. Titration of identical concentrations of NaCl showed no increase in activation at any condition, demonstrating that the riboswitch is highly specific for fluoride (**Figure 1C**, grey line), as has been previously observed from experiments performed in *E. coli* (*10*). Thus, without any optimization of riboswitch structure or function, the sensor can discriminate health-relevant concentrations of fluoride dosed into laboratory water samples.

### Changing reporters to tune sensor speed and detection threshold

Biosensor field deployment requires an output that can be quickly read with minimal supplemental equipment. Using the maximally activating fluoride concentration (3.5 mM), reactions achieved measurable signal above the OFF state in 30 minutes at 30°C, with overall 20-fold induction relative to the no-fluoride condition at the end of the 8-hour experiment (**Figure 2A**). Despite this, the sensor’s ON state was not distinguishable by eye for several hours, presenting the need for a faster reporter.

**Figure 2.**
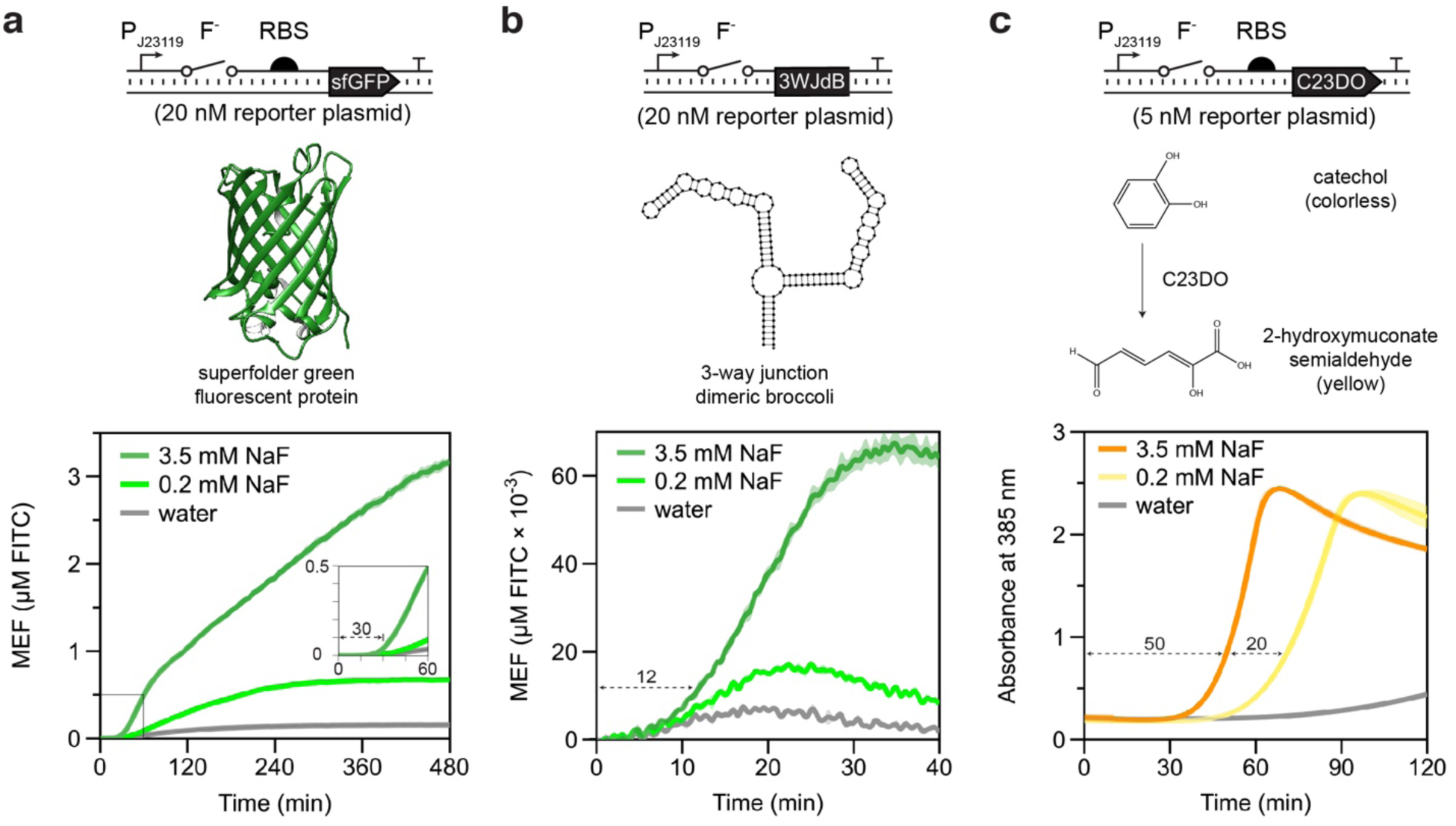
Riboswitch modularity allows fluorescent protein, RNA aptamer and enzymatic colorimetric reporter outputs. Biosensor DNA template layouts and concentrations shown above reporter information and characterization data for that reporter. **(a)** Superfolder GFP (sfGFP) reporter (structure from PDB 2B3P). Time course of reporter sfGFP induction in the presence of 3.5 mM NaF (dark green), 0.2 mM NaF (light green), or water (gray). **(b)** 3-way junction dimeric Broccoli reporter (structure predicted from NUPACK (*34*)). Time course of fluorescence in the presence of 3.5 mM NaF (dark green), 0.2 mM NaF (light green) and water (gray). **(c)** Catechol (2,3)-dioxygenase (C23DO) reporter. Reaction scheme shows the cleavage of the colorless catechol molecule into the yellow 2-hydroxymuconate semialdehyde. Time course of absorbance at 385 nm in the presence of 3.5 mM NaF (orange), 0.2 mM NaF (yellow), and water (gray). For each plot, trajectories represent average and error shading represents one standard deviation from three technical replicates. (a) and (b) are reported in mean equivalence fluorescein (MEF).

We hypothesized that we could speed sensor response with 3-way junction dimeric Broccoli (3WJdB) (*13*), an RNA aptamer that activates fluorescence of its DFHBI-1T ligand upon transcription, eliminating delays caused by translation. At all tested NaF concentrations, 3WJdB produced a signal detectable over background within 12 minutes at 30°C (**Figure 2B**), more than twice as fast as could be achieved with sfGFP (**Figure 2A**). Interestingly, this result also confirms that the fluoride riboswitch is compatible with RNA reporters, despite the potential for misfolding with the upstream riboswitch sequence. However, despite the improvement in speed, exchanging sfGFP for 3WJdB resulted in a 50-fold reduction in the sensor’s fluorescent output at the maximally activating tested condition. Thus, although the RNA-level output is preferable for its speed relative to the sfGFP output if a plate reader is accessible, it is not optimal for field deployment.

As an alternative to a fluorescent output, we used the colorimetric enzyme catechol (2,3)-dioxygenase (C23DO) as a reporter. C23DO has previously been used in genetically-encoded biosensors for plant viruses (*14*), oxidizing its colorless catechol substrate to the yellow-colored 2-hydroxymuconate semialdehyde (*15*). This color change allows gene expression to be read out either by light absorbance at 385 nanometers on a plate reader or by the visible appearance of a yellow color. All tested fluoride concentrations produced a visible output within 70 minutes at 30°C, which we empirically defined as an absorbance of 0.8 based on our previous observations (**Figure 2C)** (*14*). Notably, there was only a 20-minute time separation between the minimally and maximally activating conditions, highlighting the ability of enzymatic reporters to quickly amplify weak signals. This combination of an easily visualized output with reasonable detection speeds made C23DO our reporter of choice when designing a field-deployable diagnostic.

### Reaction tuning and lyophilization towards biosensor field deployment

We next took steps to optimize our sensor to detect fluoride near the EPA’s secondary maximum contaminant limit of 2 ppm (100 μM). We obtained a robust ON signal with our original design, but the sensor began to leak after 90 minutes (**Figure 2C**, gray line), complicating detection for trace amounts of fluoride. We attempted to mitigate this problem by reducing the amount of reporter DNA supplied to the reaction from 5 nM to 3 nM to diminish the sensor’s output. In doing so, we completely suppressed biosensor activation in the absence of fluoride (**Figure 3A**), but at the cost of significantly delaying activation.

**Figure 3.**
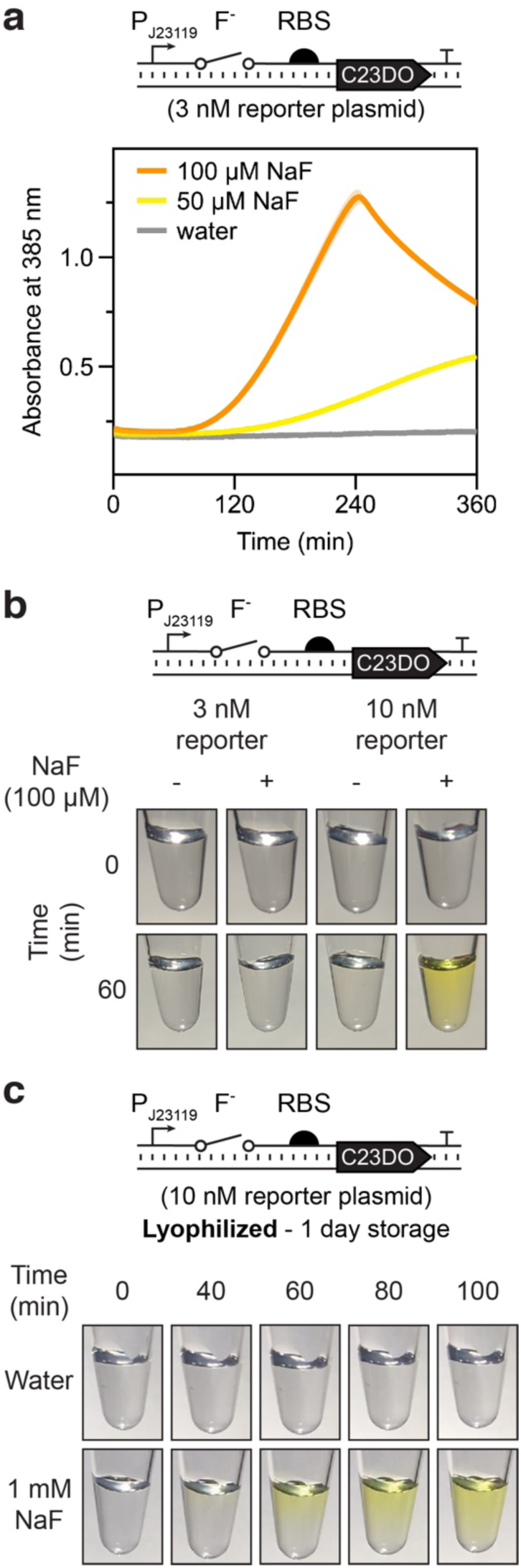
Colorimetric reporters enable fluoride sensing at environmentally relevant concentrations. **(a)** Time course of 385 nm absorbance as measured by plate reader in the presence of 100 μM NaF (orange), 50 μM NaF (yellow), and water (grey) using C23DO as a reporter. Trajectories represent average and error shading represents one standard deviation from three technical replicates. **(b)** Color change observed after 1-hour for two different reporter template concentrations with and without 100 μM NaF. Tubes were mixed by pipetting before image capture at 60 minutes. (**c)** Time lapse of rehydrated lyophilized reactions in the absence (top) and presence (bottom) of 1mM NaF.

We next hypothesized that controlling reaction timing such that the appearance of color only within a specified time window would correspond to a positive (ON) signal could improve sensitivity without sacrificing speed. To do this, we increased biosensor DNA concentration to 10 nM and increased the temperature of the CFE reaction to 37°C. Under these conditions, induction by 100 µM NaF resulted in a clear change in color within sixty minutes with no visible leak in the OFF condition (**Figure 3B, Supplemental Figure 2**). The same conditions using 3 nM DNA template resulted in no color change within 60 minutes. This result highlights an appreciable advantage afforded by the open reaction environment of cell-free systems: sensor limit of detection can be tuned simply by manipulating the reaction time and the DNA concentration of the biosensor.

Inspired by recent work that demonstrated CFE reactions can be freeze-dried and rehydrated when needed for on-demand biomanufacturing, nucleic acid detection, and educational activities (*5, 6, 16, 17*), we next aimed to demonstrate that the optimized fluoride biosensor reactions could maintain functionality after being freeze-dried. We measured the impact of lyophilization on fluoride sensitivity by flash-freezing and lyophilizing reactions containing C23DO reporter plasmid overnight. The reactions were then rehydrated with laboratory grade Milli-Q water (**Figure 3C**, top) or water containing 1 mM NaF (**Figure 3C**, bottom) and incubated at 37°C. Time-lapse photography shows visible activation within 60 minutes in the 1 mM NaF condition with no leak observed within 100 minutes in the no fluoride condition (**Supplemental Video 1**). This finding, consistent with other recent reports from cell-free systems, indicates that the fluoride riboswitch in cell-free extract is robust to lyophilization.

We also tested the viability of lyophilized reactions stored over longer periods of time. After lyophilization, reaction tubes were wrapped in Parafilm and stored in Drierite for 3 months in darkness at room temperature and atmospheric pressure before being removed and rehydrated with laboratory grade Milli-Q water or water containing 1 mM NaF. The sample rehydrated with 1 mM NaF showed strong activation within one hour, with no leak observed in the Milli-Q condition (**Supplemental Figure 3)**. Interestingly, lyophilization appeared to suppress the sensor leak in the no-fluoride condition without impacting the ability to activate expression with fluoride. The maintained viability of reactions after three months indicates that storage in desiccant and light shielding to prevent catechol oxidation are the only requirements for long-term storage of lyophilized cell-free reactions, a crucial step towards field-deployment.

### Point-of-use detection of environmental fluoride with a freeze-dried biosensor

Of particular concern in the transition from the lab to the field are components of environmental water samples, particularly natural ions like sodium, magnesium, or potassium, that could poison cell-free reactions upon rehydration. To test the robustness of our system against these sample matrix effects, we created mock fluoride-containing field samples by sampling water from a municipal tap, Lake Michigan, and an outdoor swimming pool, with Milli-Q water used as a control. NaF was then added to each sample to a final concentration of 1 mM. The biosensing reactions were prepared as before and pipetted into PCR tubes (**Figure 4A**, top) or spotted on BSA-treated chromatography paper (**Figure 4A**, bottom) before being flash frozen and lyophilized overnight. After lyophilization, reactions were immediately rehydrated with either unaltered mock field sample (-condition) or mock field sample containing 1 mM NaF (+ condition) and incubated at 37°C for one hour. For all fluoride-containing samples both in tubes and on paper, a color change was observed within one hour, with no color development in any of the no fluoride conditions. These results validate that the fluoride biosensor is robust against the unfiltered environmental samples tested and can be used in real-world conditions.

**Figure 4.**
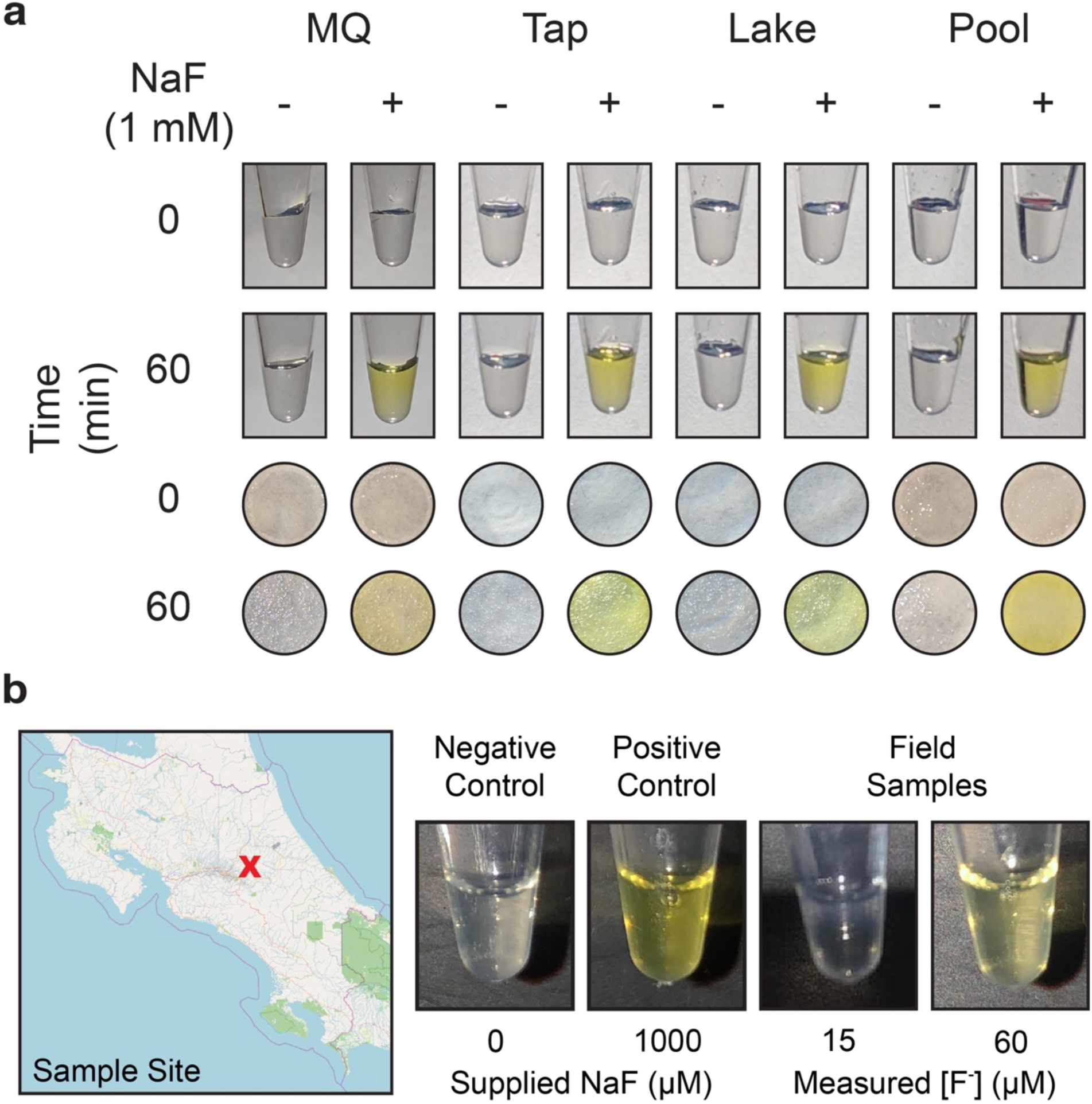
The cell-free fluoride riboswitch biosensor functions with real-world water samples and is not impacted by long-term storage and distribution. **(a)** Cell-free reactions rehydrated with various water samples with or without 1 mM NaF added. Lyophilized reactions in tubes are shown above lyophilized reactions on chromatography paper before and after one-hour incubation at 37°C. MQ = laboratory grade Milli-Q water; Tap = tap water; Lake = unfiltered Lake Michigan water; Pool = unfiltered outdoor pool water. Uncropped photos of chromatography paper experiments in **Supplemental Figure 8**) **(b)** Field testing of lyophilized cell-free reactions rehydrated with water sampled from Cartago, Costa Rica. Geographical map from OpenStreetMap (*35*). The positive control contained 1 mM NaF in the reaction before freeze-drying. The negative control was rehydrated with Milli-Q water, and the positive control and each test were rehydrated with 20 μL of unprocessed field sample. Measured fluoride concentrations obtained using a fluoride sensing electrode. Field samples are from locations B and E in **Supplementary Table 2**.

As the culmination of our optimization process, we tested our sensor’s ability to accurately classify fluoride-containing samples in the field. We specifically sought to follow a publicly available previously published environmental fluoride study that used conventional methods to sample and test publicly available natural and municipal water sources near the Irazu volcano in Cartago, Costa Rica, an area shown to have elevated fluoride levels (**Supplemental Figure 4**) (*18*). To do this, we manufactured freeze-dried fluoride biosensor reactions and transported them to Costa Rica using our simplified desiccant packaging (**Supplemental Figure 5A**) for field testing. Using sampling regions from the previous study (*18*), we collected samples in 50 mL conical tubes and tested for fluoride in batch by adding unprocessed water to a lyophilized reaction in a PCR tube with a single-use exact volume transfer pipette (**Supplemental Figure 5B**).

All field-testing was done onsite in Costa Rica without access to laboratory resources or equipment. Reactions were incubated at approximately 37°C by being held in the armpit, with reaction time increased to 5 hours to control for delayed activation from the imprecision of body heat incubation and low environmental fluoride concentrations (*14*). A strong yellow color developed in every positive control reaction within an hour, confirming robustness to reaction poisoning by potential sample matrix effects (**Supplemental Table 2**). No activation was observed within 5 hours in any samples with fluoride concentrations less than 50 μM (~1 ppm) as measured in cross-validation with a commercial fluoride-sensing electrode. However, a visible color change was observed after 3.5 hours in a water sample collected from a roadside ditch measured to have a fluoride concentration of 60 µM (**Figure 4B**). For all samples, the commercial electrode measurement confirmed the conclusions drawn from the cell-free sensors, with no false positives or false negatives observed under any conditions (n = 9) (**Supplemental Table 2**). By accurately detecting levels of fluoride relevant to public health concern thresholds in a real-world water source with minimal supplementary equipment, we have shown that the fluoride riboswitch can be effectively used as a low-cost, point-of-use cell-free diagnostic, demonstrating the potential of engineered biosensor elements for small molecule detection in the field (*19*).

## Discussion

In this work, we have demonstrated that a fluoride riboswitch can be implemented in a cell-free gene expression system to act as a field-deployable diagnostic for environmental water samples. To the best of our knowledge, this is the first demonstration of a cell-free riboswitch-based biosensor that can detect health-relevant small molecules at regulatory levels within the field. Importantly, this work represents a significant improvement in efficacy over commercially available kits (**Supplemental Figure 6)** and provides significant simplification and cost savings over gold standard electrochemical methods of fluoride detection, which cost hundreds to thousands of dollars and are cumbersome to use even for scientifically skilled operators. In contrast, our freeze-dried biosensors can currently be made for $0.40/reaction (*16*), only require a drop of water, and can be incubated with body heat.

A key strength of cell-free biosensing is that biochemical parameters such as cofactor and DNA concentration can be very easily tuned to reduce leak and improve dynamic range, which has been a historically difficult challenge for riboswitch engineering in cells. Furthermore, since riboswitches are *cis*-acting, only one DNA template concentration needs to be tuned per sensor, simplifying the optimization space relative to *trans*-acting RNA or protein regulators. When bringing these reactions to the field, we found that reactions lyophilized in PCR tubes had advantages over paper-based reactions, which rapidly dried out even when incubated in sealed, humidified containers. This effect was exacerbated by the longer incubation times for low analyte concentrations, variabilities in ambient temperature, and the practical difficulty of equipment-free incubation of paper sensors using body heat, making the tube format much more amenable to the challenges of field deployment.

This work also highlights the feasibility of using transcriptional riboswitch-mediated gene expression to convert weak-binding RNA aptamers into functional biosensors. We were surprised to find that the *B. cereus* crcB riboswitch activated so well in an *E. coli* cell-free lysate system, given the sophisticated nature of its folding mechanism and transcriptional readthrough observed both *in vitro* and *in vivo* (*11, 20*) Transcriptional riboswitches often show weak activation due to the short timescales of their regulatory decision-making, resulting in sensitivities that are kinetically, rather than thermodynamically limited (*21*). Coupling transcriptional riboswitches to enzymatic outputs like C23DO can amplify weak signals, since each reporter enzyme turns over multiple molecules of substrate (*22*). The combined kinetic mechanism of switching and the signal amplification afforded by a colorimetric reporter resulted in our sensor achieving a limit of detection of 50 µM, less than half of the lowest previously measured K_D_ for any fluoride aptamer (*23*). Thus, this work is a powerful example of why considering only thermodynamic binding affinities during aptamer selection can exclude promising, diagnostically relevant sensors.

The strategies we present here could be applied to optimize the performance of a large number of natural riboswitches for the detection of metabolites and ions relevant to environmental and human health monitoring (*24*). Additionally, the compatibility of CFE reactions for high-throughput screening (*25*) and the simple format of our DNA expression construct could be used to characterize the thousands of “orphan” riboswitches that have been bioinformatically identified but bind to unknown ligands (*26*). We imagine that these strategies could even be used to re-engineer riboswitches to have novel function (*27*–*29*). As the rules of riboswitch mechanisms are deciphered at deeper levels (*11, 30*–*32*), we hope to reach a sufficient understanding to design their functional properties to meet the global needs for field-deployable environmental and health diagnostics.

## Materials and Methods

### Plasmid Construction

Plasmids were assembled using Gibson assembly (New England Biolabs, Cat#E2611S) and purified using a Qiagen QIAfilter Midiprep Kit (QIAGEN, Cat#12143). pJBL7025 and pJBL7026 were assembled from pJBL3752. A table of all plasmid sequences can be found in Supplemental Table 1.

### Extract Preparation

Extracts were prepared according to published protocols using sonication and postlysis processing in the Rosetta2 (DE3) pLysS strain (*7*).

### CFE Experiment

CFE reactions were prepared according to established protocols (*7*). Magnesium glutamate concentration in the final reaction was optimized by extract (20 mM for shelf stability and field deployment experiments, 12 mM for all other data). Little variability was seen in extract performance between batches with varying optimal magnesium concentrations (**Supplemental Figure 7**). All kinetic CFE reactions were prepared on ice in triplicate at the 10 μL scale. 33 μL of a mixture containing the desired reaction components was prepared and then 10 μL was pipetted into three wells of a 384-well plate (Corning, 3712), taking care to avoid bubbles. Plates were sealed (ThermoScientific, 232701) and kinetic data was monitored on a BioTek Synergy H1m plate reader for sfGFP (20nM reporter plasmid, emission/excitation: 485/520 nm every five minutes for 8 hours at 30°C), C23DO (variable reporter plasmid concentration, 385 nm absorbance every 30 seconds for 4-6 hours at 30°C), and 3WJdB (20 nM reporter plasmid, emission/excitation 472/507 nm every 30 seconds for 2 hours at 30°C). C23DO reactions were supplemented with 1mM catechol and 3WJdB reactions were supplemented with 20 μM DFHBI-1T. For all fluorescence experiments, a no-DNA negative control was prepared in triplicate for every extract being tested. All reported fluorescence values have been baseline-subtracted by the average of three samples from the no-DNA condition. sfGFP and 3WJdB Measurements were then calibrated to a standard curve of fluorescein isothiocyanate (FITC) fluorescence to give standardized fluorescence units of µM equivalent FITC following a previously established procedure (*33*). Baseline subtraction was not performed for catechol reactions because reaction progress is determined from time to activation rather than maximal absorbance value. For the data depicted in Figure 1C, NaF and NaCl titrations were performed in separate experiments.

### Lyophilization

All lyophilization was performed in a Labconco FreeZone 2.5 Liter −84°C Benchtop Freeze Dryer (Cat# 710201000). A CFE reaction master mix was prepared and split into 20 μL aliquots in PCR strip tubes. Tube caps were then pierced with a pin and strips were wrapped in aluminum foil before being flash frozen in liquid nitrogen and lyophilized overnight at 0.04 mbar. After lyophilization, pierced PCR strip tube caps were replaced. Tubes were then sealed with parafilm and placed directly into Drierite (Cat#11001) for storage at room temperature (**Supplemental Figure 5A)**.

### Paper Sensors

Individual sensors were punched out of Whatman 1 CHR chromatography paper (3001-861) using a Swingline Commercial Desktop Punch (A7074020). Tickets were then placed in a petri dish and immersed in 4% BSA for one hour before being transferred to a new dish and left to air dry overnight. After drying, tickets were spotted with 20 μL of CFE reaction and placed in plastic jars (QOSMEDIX 29258), which were loosely capped and wrapped in aluminum foil before being flash frozen in liquid nitrogen and lyophilized overnight at 0.04 mbar. For testing, tickets were transferred to new jars and rehydrated with 20 μL of sample solution. Jars were then closed and sealed with parafilm before incubation for one hour at 37°C.

### Field Deployment

20 μL lyophilized reactions were prepared with 10 nM pJBL7025 and 1 mM catechol. As a positive control, additional reactions were lyophilized after being pre-enriched with 1 mM NaF. Supplementary Table 2 contains a complete list of sample site locations and water sources tested. 50 mL water samples were collected and stored in Falcon tubes (Fisher Scientific, Cat# 14-432-22) without any processing or filtration. Reactions were rehydrated by using 20 μL exact volume transfer pipettes (Thomas Scientific, 1207F80) to pull from collected samples. Three reactions were run at each sample site: (1) a positive control rehydrated with the sample, (2) a blank reaction rehydrated with the sample, and (3) a negative control reaction rehydrated with purified water to test for any reaction leak. Reactions were placed in a plastic bag and incubated at body temperature in the armpit for five hours using established protocols and marked as activated if a visible yellow color was observed (*14*). Quantitative measurements of fluoride concentration were taken with an Extech ExStik Waterproof Fluoride Meter (Cat# FL700).

## Acknowledgements

We would like to thank Professor Ana Gabriela Calderón Cornejo (Universidad de Costa Rica) and Eduardo Quirós Morales for assistance with biosensor field-testing. We also thank Ashty Karim (Northwestern University) and Professor Robert Batey (University of Colorado, Boulder) for helpful comments in preparing the manuscript, along with Khalid Alam (Stemloop, Inc.) for editing the supplemental video and Thomas Shahady (University of Lynchburg) for helpful comments about water sampling in Costa Rica. This work was supported by the Air Force Research Laboratory Center of Excellence for Advanced Bioprogrammable Nanomaterials (C-ABN) Grant FA8650-15-2-5518 (to M.C.J. and J.B.L), the David and Lucile Packard Foundation (to M.C.J.), an NSF CAREER Award (1452441 to J.B.L.), and the Camille Dreyfus Teacher-Scholar Program (to M.C.J. and J.B.L.). A.D.S. was supported in part by the National Institutes of Health Training Grant (T32GM008449) through Northwestern University’s Biotechnology Training Program. The views and conclusions contained herein are those of the authors and should not be interpreted as necessarily representing the official policies or endorsements, either expressed or implied, of the Air Force Research Laboratory, Air Force Office of Scientific Research, or US Government.

## Author Contributions

Conceptualization, W.T., A.D.S., M.C.J., & J.B.L.; Data Curation, W.T., A.D.S., & M.S.V.; Formal Analysis, W.T & A.D.S.; Investigation, W.T., A.D.S, & M.S.V.; Methodology, W.T., A.D.S., M.S.V., & J.B.L.; Project administration, W.T. & J.B.L.; Validation, W.T., A.D.S., & M.S.V.; Funding acquisition, N.K., M.C.J., & J.B.L.; Writing – original draft, W.T., A.D.S., & J.B.L.; Writing – review & editing, W.T., A.D.S., M.S.V., M.C.J., & J.B.L.

## Competing Interests Statement

The authors have submitted one provisional patent application (U.S. Patent Application Serial No. 62/813,368) for the technologically important developments included in this work. J.B.L is a cofounder of Stemloop, Inc. J.B.L.’s interests are reviewed and managed by Northwestern University in accordance with their conflict of interest policies.

## Data Availability

All source data for main and SI figures was deposited open access in Northwestern’s Arch database (https://arch.library.northwestern.edu). Data can be accessed via https://doi.org/10.21985/N2RJ64.

## Plasmid Availability

All plasmids are being deposited in Addgene with accession numbers 128809 – 128811.

## Notes

https://arch.library.northwestern.edu/concern/generic_works/5q47rn983?locale=en

