## Supplementary Information for "Point-of-Use Detection of Environmental Fluoride via a Cell-Free Riboswitch-Based Biosensor"

*Short Title: Point-of-Use Fluoride Biosensing*

Walter Thavarajah[1,3,4], Adam D. Silverman[1,3,4], Matthew S. Verosloff[2,3,4], Nancy Kelley-Loughnane[5], Michael C. Jewett[1,3], and Julius B. Lucks[1,3,4]

1 - Department of Chemical and Biological Engineering, Northwestern University, 2145 Sheridan Rd, Evanston, IL, 60208, USA
2 - Interdisciplinary Biological Sciences Graduate Program, Northwestern University, 2204 Tech Drive, Evanston, IL, 60208, USA
3 - Center for Synthetic Biology, Northwestern University, 2145 Sheridan Rd, Evanston, IL, 60208, USA
4 - Center for Water Research, Northwestern University, 2145 Sheridan Rd, Evanston, IL, 60208, USA
5 - Materials and Manufacturing Directorate, Air Force Research Laboratory, Wright-Patterson Air Force Base, Ohio 45433, United States

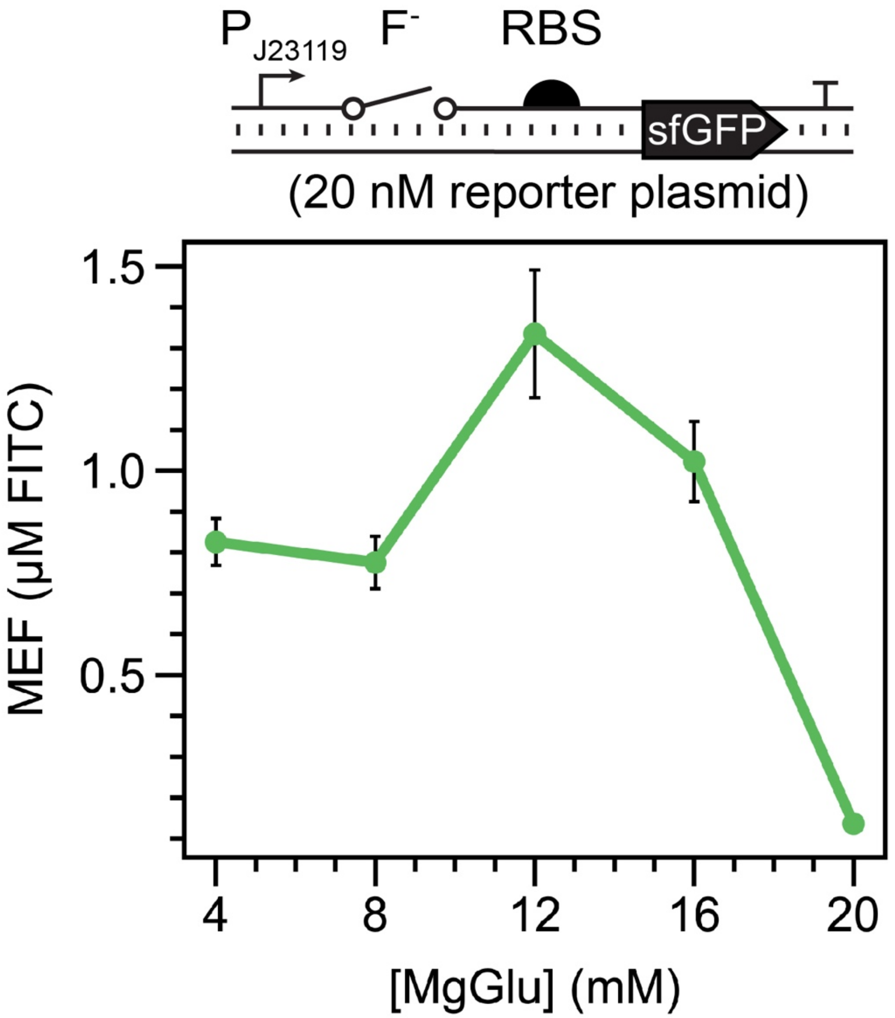


**Supplemental Figure 1. Fluoride riboswitch magnesium optimization.** Reactions were supplied with 1 mM NaF and varying magnesium glutamate as indicated. Data shown are endpoint measurements from an eight-hour experiment. Error bars represent one standard deviation from three technical replicates. This experiment indicated that a magnesium glutamate concentration of 12 mM gave optimal fluorescence, though we note that optimal magnesium concentration can vary between extracts.

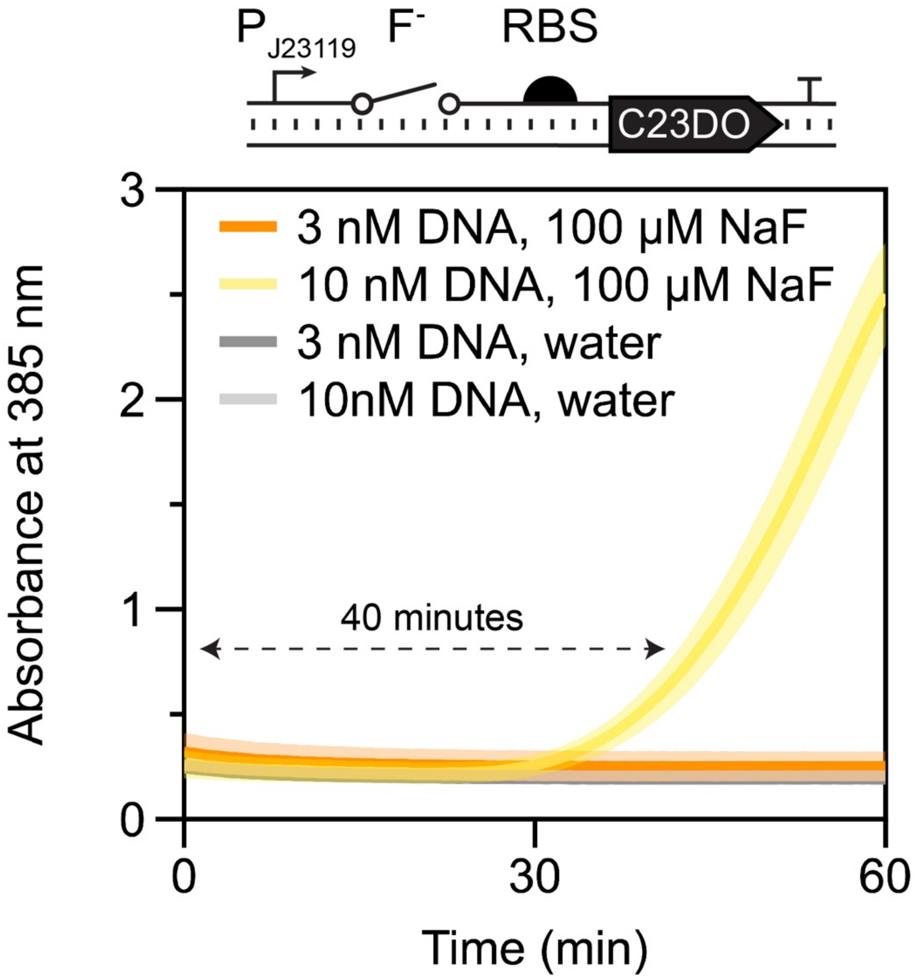


**Supplemental Figure 2. Kinetic traces for reaction conditions depicted in Figure 3B.** This experiment was run at 37°C to best mirror experimental conditions for reactions run in PCR tubes. Visible activation is seen in 40 minutes for the condition with 10 nM biosensor DNA and 100 µM NaF, corroborating the results depicted in Figure 3B. Trajectories represent average and error shading represents one standard deviation from three technical replicates

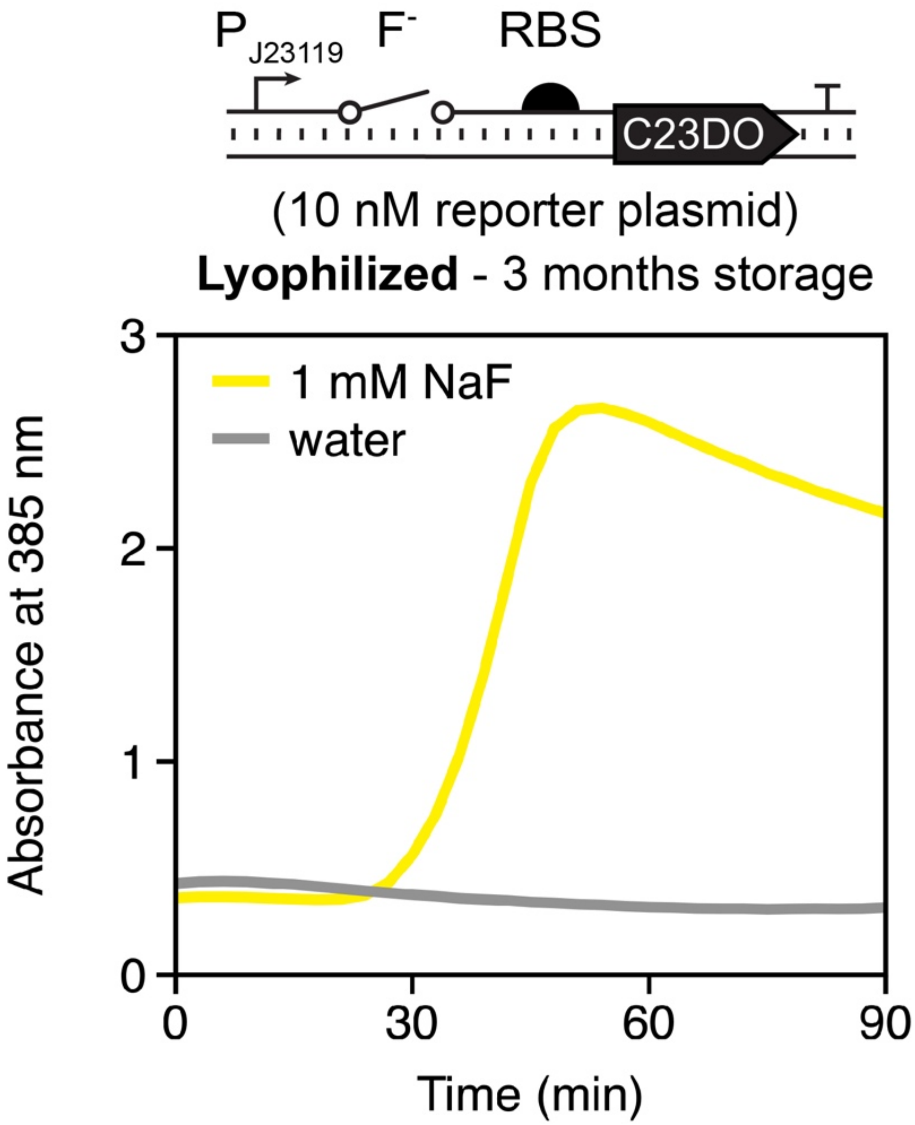


**Supplemental Figure 3. Lyophilized reactions remain viable after three months of storage in desiccant.** Reactions were stored in darkness under ambient conditions before rehydration with 20μL of water with or without 1mM NaF. Trajectories represent data from one experiment.

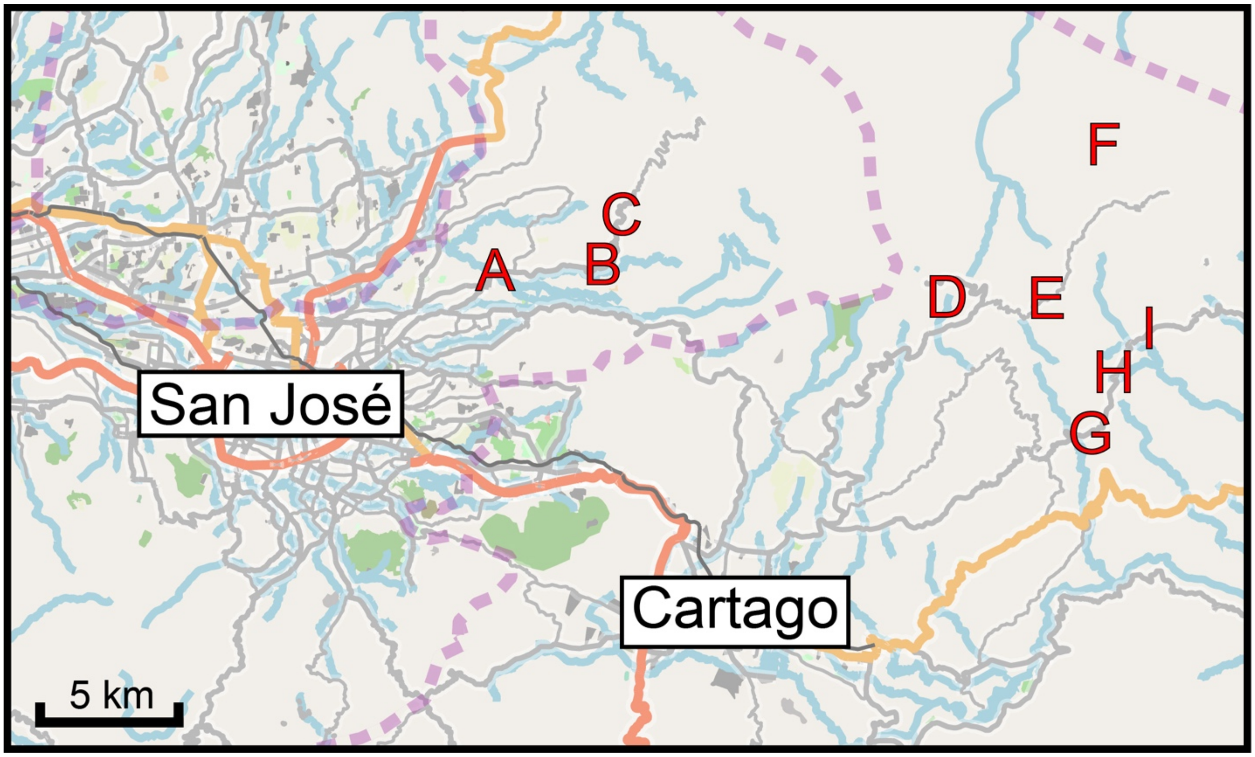


**Supplemental Figure 4. Map of Costa Rican water sampling locations.** Sampling locations were determined from a previously published report about the presence of fluoride in water within this region (*1*). Each letter represents a unique source where 50 ml of water was sampled. Locations center around the Irazu volcano, a known source of fluoridated salts (*1*). Data presented in Figure 4B is from location E.

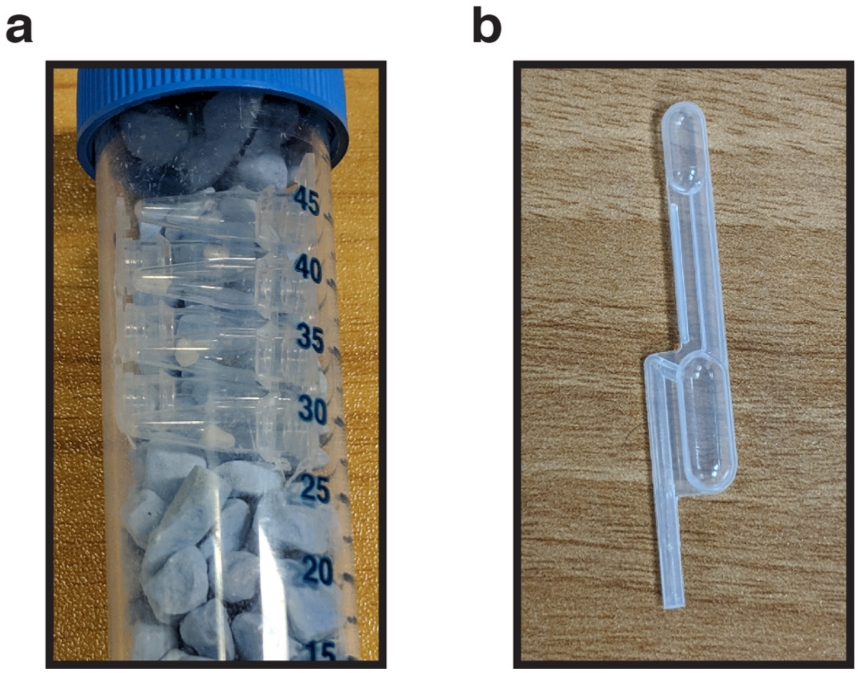


**Supplemental Figure 5. Necessary field-testing equipment. (a)** Lyophilized reactions stored in a 50 mL conical tube filled with desiccant. Reactions can be individually removed from the strip for testing on demand. Because the reactions are not stored under nitrogen gas, the tube can be opened and resealed as many times as necessary. **(b)** 20 μL exact volume transfer pipette (Thomas Scientific, 1207F80). Pipettes measure approximately 5 cm lengthwise. By squeezing and releasing the bulb on top, 20 μL of fluid is transferred into the stem, with any excess entering the overflow reservoir. Squeezing the bulb again dispenses the water, which is added directly to the lyophilized reactions before incubation.

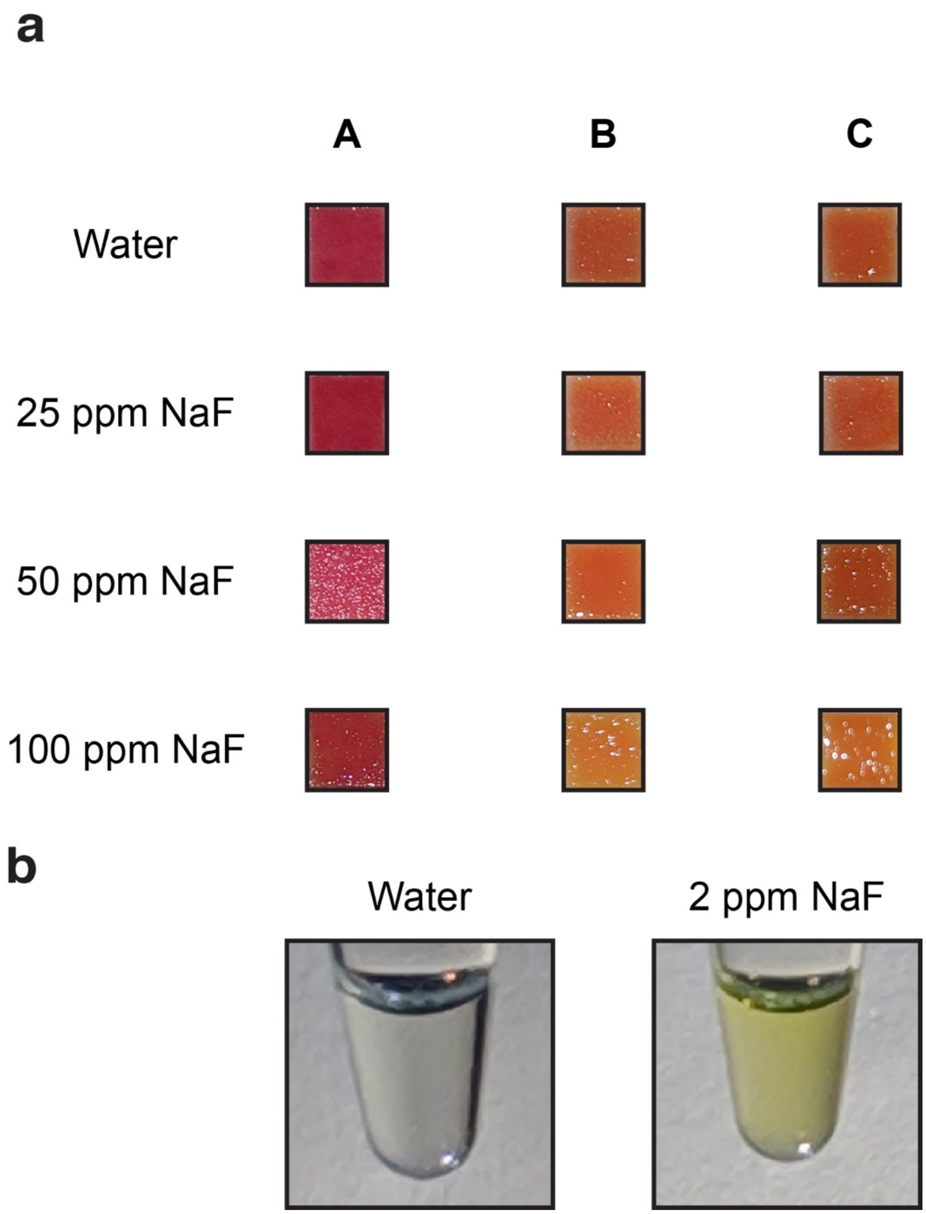


**Supplemental Figure 6. The cell-free fluoride riboswitch biosensor is capable of higher-fidelity sensing than currently available colorimetric assays. (a)** Color change from three anonymized commercially available test strips with a reported sensitivity range between 10-100+ ppm of fluoride. Strips were dipped in the indicated NaF solution and held at room temperature for 30 seconds to wait for color change, as directed by supplied instructions. No readily apparent change was observed at any fluoride concentration. **(b)** Fluoride detection using a cell-free reaction containing 10 nM of the fluoride riboswitch regulated C23DO DNA template. The reaction was set up, incubated at 37°C for 1 hour, then mixed by pipetting before image capture. Despite the delayed activation due to time consumed by transcription and translation, clear activation can be seen at concentrations below 10 ppm.

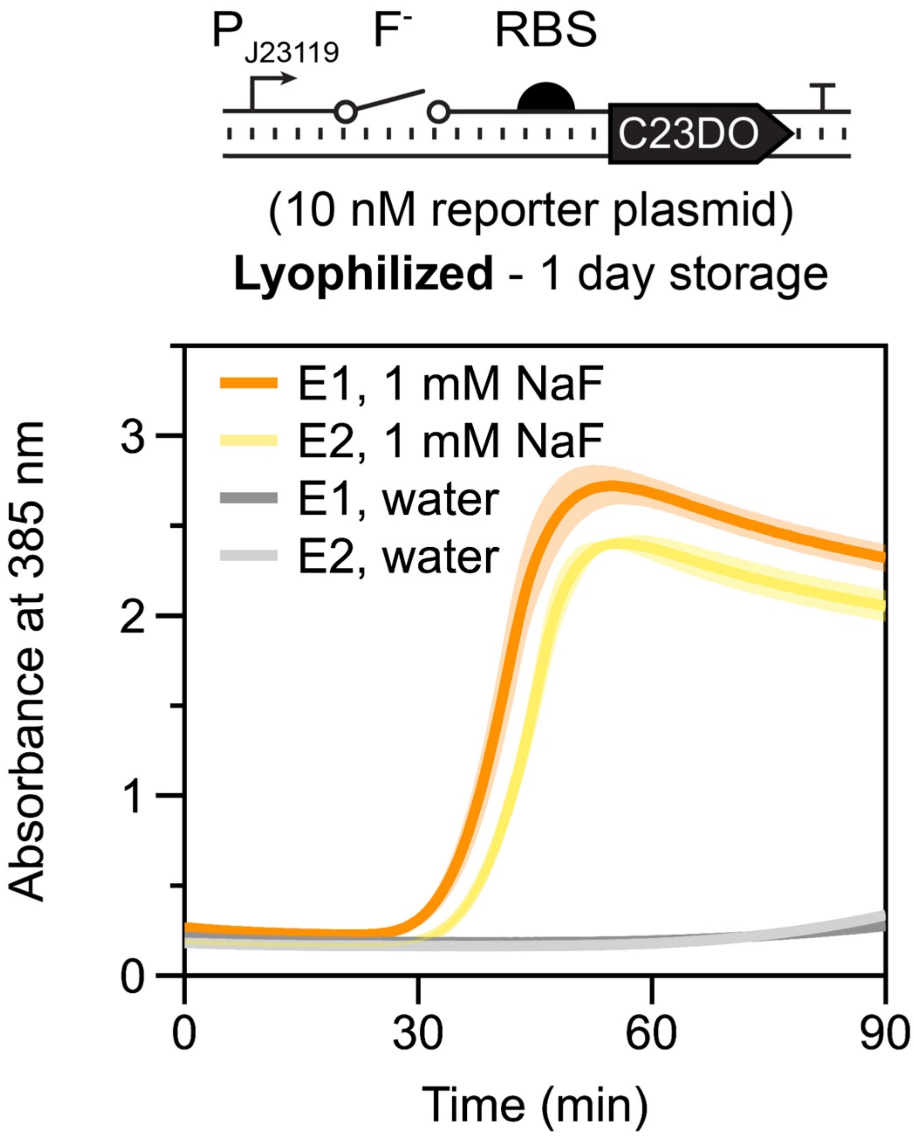


**Supplemental Figure 7. Lyophilized reactions show little variability between batches of cell-free extract.** Cell-free reactions containing fluoride riboswitch-regulated C23DO were set up with different batches of cell-free extract (E1 and E2) and lyophilized overnight. The next morning, reactions were rehydrated and reaction progress, read out by absorbance at 385 nm, was monitored in a plate reader maintained at 30°C. The reactions reached maximal activation almost simultaneously in both conditions containing 1mM NaF (orange and yellow lines) and had negligible leak without added NaF (gray and dark gray lines).

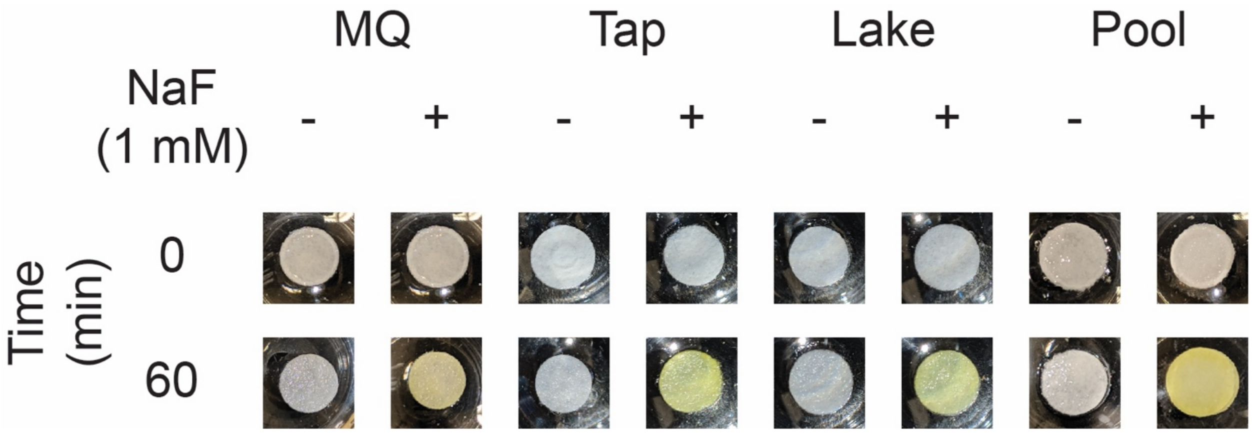


**Supplemental Figure 8. Uncropped photos of lyophilized cell-free reactions on paper.** Reactions pictured here are the same as those pictured in Figure 4A.

**Supplemental Table 1. Sequences of constructs.** Constructs utilized the Anderson promoter BBa_J23119_Spe1 (blue), the B. cereus fluoride riboswitch (red), a ribosome binding site (RBS) (pink), superfolder GFP (sfGFP) (green), catacholase 2,3 dioxygenase (C23DO) (yellow), three-way junction dimeric broccoli (3wjdb) (teal), and T1/TE terminator (grey).

| **Name** | **Sequence 5' to 3'** |
|---|---|
| **pJBL3752:**<br>Anderson Promoter BBa_J23119_Spe1 / B.cereus Fluoride Riboswitch / RBS / Superfolder GFP coding sequence/ T1/TE terminator | ttgacagctagctcagtcctaggtataatactagtttaTAGGCGATGGAGTTCGCCATA<br>AACGCTGCTTAGCTAATGACTCCTACCAGTATCACTACTGGTAGGA<br>GTCTATTTTTTTaggaggaaggatctatgagcaaaggagaagaacttttcactggagttgt<br>cccaattcttgttgaattagatggtgatgttaatgggcacaaattttctgtccgtggagagggtgaagg<br>tgatgctacaaacggaaaactcacccttaaatttatttgcactactggaaaactacctgttccgtggc<br>caacacttgtcactactctgacctatggtgttcaatgcttttcccgttatccggatcacatgaaacggca<br>tgactttttcaagagtgccatgcccgaaggttatgtacaggaacgcactatatctttcaaagatgacg<br>ggacctacaagacgcgtgctgaagtcaagtttgaaggtgataccttgttaatcgtatcgagttaaa<br>gggtattgattttaaagaagatggaaacattcttggacacaaactcgagtacaactttaactcacac<br>aatgtatacatcacggcagacaaacaaaagaatggaatcaaagctaacttcaaaattcgccaca<br>acgttgaagatggttccgttcaactagcagaccattatcaacaaaatactccaattggcgatggccc<br>tgtccttttaccagacaaccattacctgtcgacacaatctgtcctttcgaaagatcccaacgaaaagc<br>gtgaccacatggtccttcttgagtttgtaactgctgctgggattacacatggcatggatgagctctaca<br>aataaggatccaaactcgagtaaggatctccaggcatcaaataaaacgaaaggctcagtcgaa<br>agactgggcctttcgttttatctgttgtttgtcggtgaacgctctctactagagtcacactggctcaccttc<br>gggtgggcctttctgcgtttata |
| **pJBL7025:**<br>Anderson Promoter BBa_J23119_Spe1 / B.cereus Fluoride Riboswitch / RBS / Catechol 2,3-dioxygenase coding sequence/ T1/TE terminator | ttgacagctagctcagtcctaggtataatactagtttaTAGGCGATGGAGTTCGCCATA<br>AACGCTGCTTAGCTAATGACTCCTACCAGTATCACTACTGGTAGGA<br>GTCTATTTTTTTaggaggaaggatctatgaacaaaggtgtaatgcgaccgggccatgtgc<br>agctgcgtgtactggacatgagcaaggccctggaacactacgtcgagttgctgggcctgatcgag<br>atggaccgtgacgaccagggccgtgtctatctgaaggcttggaccgaagtggataagttttccctgg<br>tgctacgcgaggctgacgagccgggcatggattttatgggtttcaaggttgtggatgaggatgctctc<br>cggcaactggagcgggatctgatggcatatggctgtgccgttgagcagctacccgcaggtgaact<br>gaacagttgtggccggcgcgtgcgcttccaggccccctccgggcatcacttcgagttgtatgcaga<br>caaggaatatactggaaagtgggggtttgaatgacgtcaatcccgaggcatggccgcgcgatctga<br>aaggtatggcggctgtgcgtttcgaccacgccctcatgtatggcgacgaattgccggcgacctatg<br>acctgttcaccaaggtgctcggtttctatctggccgaacaggtgctggacgaaaatggcacgcgcg<br>tcgcccagtttctcagtctgtcgaccaaggcccacgacgtggccttcattcaccatccggaaaaag<br>gccgcctccatcatgtgtccttccacctcgaaacctgggaagacttgcttcgcgccgccgacctgat<br>ctccatgaccgacacatctatcgatatcggcccaacccgccacggcctcactcacggcaagacc<br>atctacttcttcgacccgtccggtaaccgcaacgaagtgttctgcgggggagattacaactacccgg<br>accacaaaccggtgacctggaccaccgaccagctgggcaaggcgatcttttaccacgaccgcat<br>tctcaacgaacgattcatgaccgtgctgacctgataaggatccaaactcgagtaaggatctccagg<br>catcaaataaaacgaaaggctcagtcgaaagactgggcctttcgttttatctgttgtttgtcggtgaac<br>gctctctactagagtcacactggctcaccttcgggtgggcctttctgcgtttata |

| | |
|---|---|
| **pJBL7026:** Anderson Promoter BBa_J23119_Spe1 / B.cereus Fluoride Riboswitch / 3-Way Junction Dimeric Broccoli coding sequence / T1/TE terminator | ttgacagctagctcagtcctaggtataatactagtttaTAGGCGATGGAGTTCGCCATAAACGCTGCTTAGCTAATGACTCCTACCAGTATCACTACTGGTAGGAGTCTATTTTTTTcccacatactctgatgatccgagacggtcgggtccagatattcgtatctgtcgagtagagtgtgggctcggatcattcatggcaagagacggtcgggtccagatattcgtatctgtcgagtagagtgtgggctcttgccatgtgtatgtgggccaggcatcaaataaaacgaaaggctcagtcgaaagactgggcctttcgttttatctgttgtttgtcggtgaacgctctctactagagtcacactggctcaccttcgggtgggcctttctgcgtttata |

**Supplemental Table 2. GPS coordinates and documentation for water sampling sites depicted in Supplemental Figure 4. GPS coordinates are reported to the nearest ten minute resolution and thus represent regions sampled rather than exact locations.** Measured concentrations were determined with a fluoride sensing electrode. “Activation” refers to the production of a visually detectable yellow color after sensor rehydration (see Methods). Data presented in Figure 4B is from location E. Permissions were received before sampling indoor faucets.

| Site | GPS Coordinates | Source | Measured $[F^-]$ (ppm) | Activation | Negative Control | Positive Control |
|---|---|---|---|---|---|---|
| **A** | 9°58'30"N 84°00'20"W | Indoor Faucet | 0.2 | No | Off | On |
| **B** | 10°00'00"N 83°57'30"W | Muddy Ditch | 0.3 | No | Off | On |
| **C** | 10°00'40"N 83°57'10"W | Indoor Faucet | 0.2 | No | Off | On |
| **D** | 9°56'50"N 83°51'50"W | Outdoor Supply | 1 | Yes | Off | On |
| **E** | 9°58'10"N 83°49'40"W | Muddy Ditch | 1.2 | Yes | Off | On |
| **F** | 10°00'10"N 83°46'50"W | Outdoor Supply | 0.5 | No | Off | On |
| **G** | 9°56'30"N 83°46'40"W | River | 0.1 | No | Off | On |
| **H** | 9°57'10"N 83°46'20"W | River | 0.3 | No | Off | On |
| **I** | 9°57'20"N 83°46'20"W | River | 0.3 | No | Off | On |

**Supplemental Video 1. Time lapse of sensor activation depicted in Figure 3C.** Tubes were rehydrated with either water (left) or 1 mM NaF (right). Total time elapsed is 100 minutes.
